# Chromatin is frequently unknotted at the megabase scale

**DOI:** 10.1101/762872

**Authors:** Dimos Goundaroulis, Erez Lieberman Aiden, Andrzej Stasiak

## Abstract

Knots in the human genome would greatly impact diverse cellular processes ranging from transcription to gene regulation. To date, it has not been possible to directly examine the genome *in vivo* for the presence of knots. Recently, methods for serial fluorescent *in situ* hybridization have made it possible to measure the 3d position of dozens of consecutive genomic loci, in vivo. However, the determination of whether genomic trajectories are knotted remains challenging, because small errors in the localization of a single locus can transform an unknotted trajectory into a highly-knotted trajectory, and vice versa. Here, we use stochastic closure analysis to determine whether a genomic trajectory is knotted in the setting of experimental noise. We analyse 4727 deposited genomic trajectories of a 2Mb long chromatin interval from chromosome 21. For 243 of these trajectories, their knottedness could be reliably determined despite the possibility of localization errors. Strikingly, in each of these 243 cases, the trajectory was unknotted. We note a potential source of bias, insofar as knotted contours may be more difficult to reliably resolve. Nevertheless, our data is consistent with a model where, at the scales probed, the human genome is often free of knots.

## Introduction

When freshly divided eukaryotic cells, such as human cells, enter the interphase stage of the cell cycle their chromosomes decondense and spread within the volume of chromosomal territories (1). Each decondensed chromosome contains one very long chromatin fibre, which in average-sized human chromosome contains about 130 million of base pairs (http://www.ensembl.org/Homo_sapiens/Location/Genome?r=Y:1-1000). Since the linear density of chromatin fibres is of about 100 bp/nm (2), the length of chromatin fibre of an average chromosome is of about 1300 micrometers. This length is sufficient to traverse several hundred times across Individual chromosome territories, which are roughly spherical and have a diameter is of about 2 micrometers (1). Chromatin fibres in chromosomal territories are associated with DNA topoisomerases that through transient cleavages and religations can permit interacting regions of chromatin fibres to pass through each other (3,4). Given these considerations, the natural expectation was that chromatin fibres in interphase chromosomes form very complex knots (5). However, studies measuring how the probability of contacts between chromosomal loci decreases with the genomic distance separating these loci, excluded the possibility that decondensed chromosomes form such complex knots as could be formed if chromatin fibres could freely pass through each other in topoisomerase-mediated reactions (5). The observed scaling profiles relating contact probability to the genomic distance did not exclude though the possibility that relatively simple chromatin knots can be scattered within decondensed chromosomes (6).

Due to linear structure of eukaryotic chromosomes, such unequivocal methods of detection of DNA knots on circular DNA molecules as gel electrophoresis (7-9) or electron microscopy (10,11) can’t be used to directly answer the question of whether interphase human chromosomes are knotted or not. Gel electrophoresis can be though used to study formation of knots in circular yeast minichromosomes, which are naturally existing, autonomously replicating, extrachromosomal genetic elements with normal chromatin structure (9). Very recent studies of yeast minichromosomes revealed that as the size of studied yeast minichromosomes increases from 1.4 kb to 11.7 kb, the steady state fraction of knots initially increases with the size of minichromosomes but then stabilizes at the value of about 3%, upon reaching the size of 8 kb (9). The observation that the probability of knotting of chromatin rings forming minichromosomes does not exceed 3%, irrespectively of the size of circular minichromosomes, suggests that also normal yeast chromosomes, with the average size of about 750 kb, may show similarly low level of knotting (9). The question arises though whether chromatin fibres in interphase chromosomes of higher eukaryotes, such as humans, are knotted and what is the frequency of these knots. As crowding of chromatin in cells of higher eukaryotes is stronger than in yeast cells, the equilibrium knotting level could be correspondingly higher.

On the other hand, knotting level in chromatin constituting chromosomes of higher eukaryotes, such as humans, may be much lower than this detected in circular minichromosomes of yeast. Replication of circular DNA molecules encounters particular topological difficulties (12,13) and these topological difficulties were shown to result in formation of knots in replicating DNA molecules (14-16). As chromatin fibres in human chromosomes are not circular, the process of their replication may not generate knots.

Indeed, early studies using Hi-C to measure the contact probability between pairs of loci genome-wide were consistent with such models of chromatin folding as fractal globule in which knots are absent (5) or models where knotting is very limited and may consist at most of simple knots scattered within decondensed chromosomes (6). Also, newer theoretical studies based on Hi-C data showed that block co-polymer nature of chromatin, where regions with the same epigenetic marks attract each other, suggested that the presence of knots is thermodynamically disfavoured, which in turn would direct DNA topoisomerase-mediated passages of chromatin fibres through each other towards unknotting (17). In addition, the process of chromatin loop extrusion could also provide a very efficient way to unknot chromatin (18,19).

Until recently there were no methods of structural analysis of entire chromosomes that could permit detection and characterization of individual knots formed on chromatin fibres within eukaryotic chromosomes. However, recently, such methods as single cell Hi-C opened the possibility to reconstruct chromatin paths in individual chromosomes at the moment of cell fixation (20). Previous methods of population Hi-C could only give an information about the average path of chromatin fibres constituting a given chromosome in millions of cells of a given type (5,21,22) and since chromosomes are highly dynamic (23) one could not use the population HI-C data to search for chromatin knots in individual chromosomes. Very recent studies using numerical simulations to analyse single cell Hi-C data suggested that chromatin fibres in individual chromosomes can be knotted (24-27). However, the concluded knotting level was low with just one trefoil knot that appeared consistently in independent numerical simulations starting from the same single cell Hi-C data of human chromosome 14 (27). However, the apparent paucity of chromatin knots concluded from single cell Hi-C data may be the consequence of unavoidable low resolution of single-cell Hi-C approach that currently does not exceed 100 kb (20). Therefore, in numerical simulations based on single cell Hi-C data individual chromosomes are modelled as continuous polygonal chains with each segment representing a 100 kb large portion of chromatin (24). It is obvious that if there were relatively tight chromatin knots with their average size of about 100 kb, they would not be realizable as knots in chromosomes modelled with 100 kb resolution, as they would be represented by just one or two straight segments in modelled chromosomes. It is less obvious, though, that to form even a simplest knot using a polygonal chain one needs at least six segments (28). For this reason, all possible chromatin knots which were not spread over a chromatin portion larger than 500 kb could only form unknotted portions of the polygonal chain representing a given chromosome modelled at 100 kb resolution. In fact, the reproducibly observed trefoil knot in modelling studies based on single cell Hi-C data was spread over 25 segments, which corresponds to 2.5 Mb large chromatin portion (24,27).

To probe the knottedness of chromatin at the smaller scale than these probed using single cell Hi-C approach one would need a method that can provide us with the spatial positions of chromatin intervals smaller than 100 kb. Very recently, an exciting new approach has emerged, combining Oligopaint chromatin labelling with high resolution optical imaging in order to determine the position of a series of consecutive loci (29,30). Using this approach, the centroid position of many sequential 30kb-large chromatin loci spanning a 2Mb long chromosomal region were determined with 50 nm accuracy (29).

Here, we use stochastic closure analysis to determine whether a genomic trajectory is knotted in the setting of realistic experimental noise. We use our method to look for knots in 4727 chromatin tracings generated by Bintu et al., (29) and which are publicly available at https://raw.githubusercontent.com/BogdanBintu/ChromatinImaging/master/Data/IMR90_chr21-28-30Mb.csv. We find that the true topology of the region could be inferred in 243 cases. Strikingly, in each case we found that the region was unknotted.

## Material & Methods

### Oligopaint chromatin tracing

We analyse here chromosome tracings deposited by Bintu et al. (29), The detailed description of the complex Oligopaint method of simultaneous tracing of selected genomic regions in chromosomes within thousands of fixed cells can be found in Bintu et al (29). Here we just briefly describe the principle of the method. The Oligopaint method used by Bintu et al., permitted the authors to label one after the other many consecutive 30 kb-long regions of chromatin constituting together ca 2 Mb large chromatin fragment. Each 30 kb-long chromatin region was hybridized with ca 300 fluorescent probes that were practically uniformly redistributed within each 30 kb fragment. Diffraction limited 3-D image of each labelled 30 kb fragment served to determine the X, Y and Z coordinates of the centroid position of each region. Once imaging of cells with the first labelled 30 kb fragment was finishes, the labelling was removed and the next 30 kb fragment was labelled with another 300 fluorescent probes and new images were acquired. The procedure was repeated for over 60 cycles, till the desired regions with up to 2 MB were traced. Control reactions, where a given region was labelled for the second time, showed that the two experimental localizations of centroid position of the same 30 kb-long chromatin portions differed from each other but the difference did not exceed 50 nm. Therefore, the precision/error determining spatial positions of sequential 30kb-long chromatin portions by Oligopaint method is of about 50 nm (29). The tracings of individual 2 Mb-long chromatin regions are represented by polygonal curves, which sequential vertices are the Oligopaint-determined positions of centroids of consecutive 30 kb-long chromatin regions in a given analysed chromosome.

### Numerical closure and knot type determination of individual chromosome tracings

Individual polygonal curves determined by X, Y and Z coordinates of individual tracings were closed as described earlier (31). In brief, each polygonal curve was closed 100 times by adding each time two very long segments starting from the two end points of the curve and being directed parallel to each other. The closure was completed by adding a straight segment joining the two distal ends of the two long segments. The procedure was repeated 100 times but each time the direction of both added parallel, long segments was changed. The 100 directions of added long segments were equally redistributed in the space. The knot type resulting from each closing direction was determined by calculation of Jones and HOMFLY-PT polynomials (32-34). The knot type that was most frequently identified among 100 differently closed curves was then assumed to represent the topology of a given tracing. In figures and tables, we apply the Alexander-Brigs notation of knots where the first number indicates the minimal number of crossings a given knot can show in a projection and where the second number, written as subscript, indicates the tabular position of that knot in standard tables of knots among knots with the same number of crossings (35). Thus, for example the notation 3_1_ indicates a knot type which in its minimal crossing representation shows three crossings. The subscript 1 indicates that this knot in standard tables of knots is presented at the first position among knots with three crossings, although there is only one knot type with three crossings. However, as the number of possible knot types increases with the their number of crossings, the notation 9_30_ indicates a knot type that in standard tables of knots is presented at the 30th position among 49 different prime knots which minimal crossing number is 9. In case of knots with more than 10 crossings the notations include also letters a or n to indicate whether a given knot is alternating or non-alternating, respectively (36). The notation of composite knots resulting from tying of two or more separate prime knots contains # sign between the corresponding prime knots.

### Topological analysis of all subchains in polygonal chains

As all subchains in polygonal chains are linear, their topological analysis is performed similarly to topological analysis of the entire chains. Each subchain was subject to stochastic multiple closure procedure and the knot type observed most frequently among closed chains was attributed to a given subchain. The results of topological analysis of all subchains of a given larger chains are conveniently presented as a matrix in which a colour of each cell tells us what is the knot type of the subchain that starts with the vertex indicated on the X axis and ends with the vertex indicated on the Y axis (37).

### Numerical simulation of topological consequences of the limited precision of tracing procedure

To simulate the effect of a limited precision of the determination of the centroid position of sequential 30kb-long chromatin blocks, we took coordinates of individual deposited tracings and introduced method-specific tracing errors to positions of every vertex. The new values of X, Y and Z coordinates of each vertex were taken from the normal distribution centred at the original position. The normal distribution function along each axis was scaled in such a way that its standard deviation coincided with 50 nm distance from the original position. In addition, we rejected all trial displacements where the combinations of displacements along the X, Y and Z axis resulted in a 3D displacement from the original point being larger than 50 nm. If a trial displacement was rejected a new displacement was tested. Using this procedure, for every analysed tracing, we produced 10 independent tracings, where each one was derived from the original deposited tracing. Each of error-perturbed tracings was then subject of stochastic multiple closure to determine the knot type conditioned by a given error-perturbed tracing. We analysed then whether error-induced perturbations resulted in changes of the knot type as compared to the knot type of the deposited, non-perturbed tracings.

## Results

### Visualization and analysis of deposited chromosomal tracings

Figure 1 shows a polygonal chain corresponding to the first deposited tracing (IMR90_chr21-28-30Mb.csv chr_idx:1) of ca 2Mb-long chromatin fibre constituting the region 28-30 MB of human chromosome 21, studied by Bintu et al., (29). The sequential vertices of the shown chain correspond to centroid positions of sequential 30kb long chromatin portions as determined by Oligopaint chromatin tracing. The diameter of all segments in the chain is set to correspond to the physical diameter of 10 nm as this reflects the diameter of chromatin fibres in interphase chromosomes (38). The two spherical beads placed at the ends of polygonal chains have their diameter set to correspond to a physical distance of 50 nm as this is the reported error range in the determination of the centroid positions of sequential 30 kb long chromatin regions (29). It is visible that in many places the non-consecutive segments of the chain approach each other to a distance much smaller than 50 nm. Therefore, considering the limited precision of the method, one can’t be certain whether the tracing reflects correctly the topology of the traced region. However, the shown tracing, as it is, is topologically characterized as forming 9_30_ knot (see below how the knot type of open polygonal curves is here determined). A standard, minimal crossing representation of that knot is shown in an insert in Figure 1.

**Figure 1.**
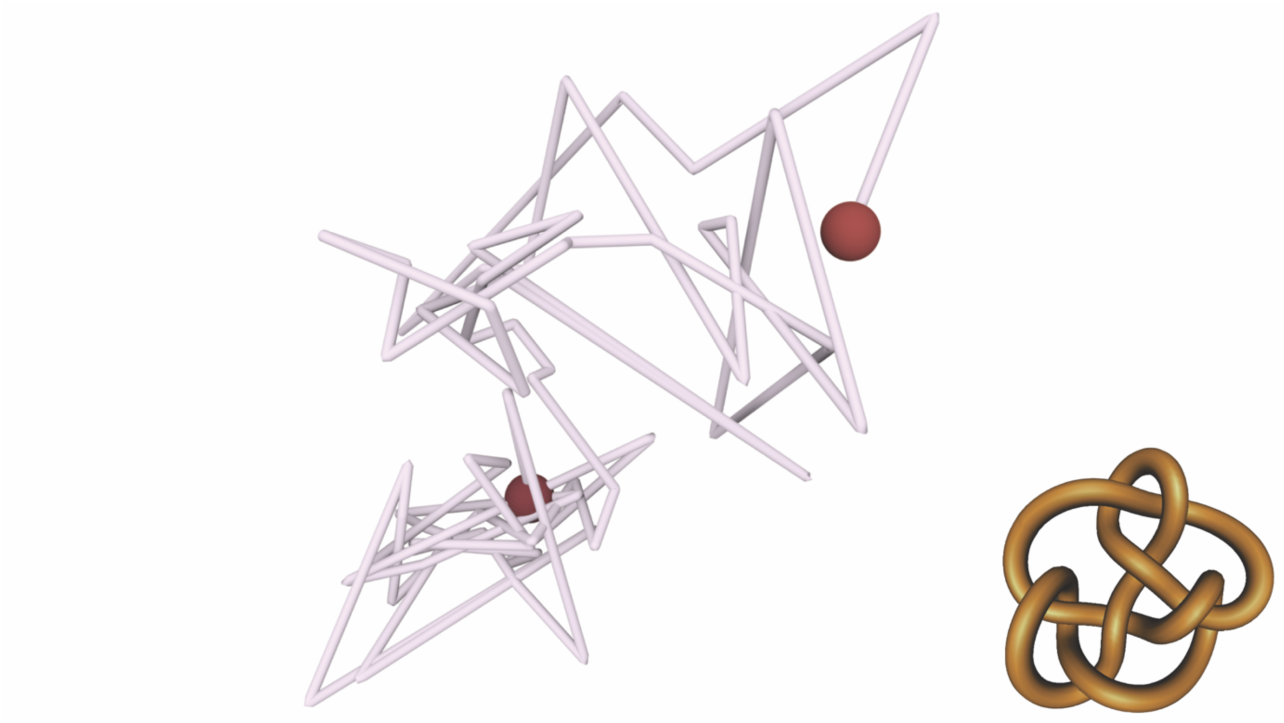
Polygonal chain determined by one of deposited chromatin tracings by Bintu et al. (29). The X, Y and Z coordinates of sequential vertices are the coordinates of centroid positions of sequential 30 kb-long chromatin portions as measured using Oligopaint method (29). The shown polygonal chain corresponds to the tracing IMR90_chr21-28-30Mb.csv chr_idx:1, which multiple closure analysis reveals that it forms 9_30_ knot. The diameter of segments is set to correspond to the physical diameter of 10 nm to reflect the diameter of 10nm chromatin fibres. The diameter of large beads, placed at both ends of the polygonal chain, is set to 50 nm to reflect the error range in the determination of the position of centroid positions of 30kb long chromatin portions using the Oligopaint method. Inset shows a minimal crossing representation of 9_30_ knot.

### Statistical analysis of the global knottedness of 4272 independently traced 2 MB large chromosomal regions

In a strict topological sense, knots can be only defined for closed paths in space. However, individual chromatin fibres forming entire chromosomes in eukaryotic cells are linear and this of course also applies to traced chromosomal regions analysed here. To be able to characterize the topology of open paths in space one needs to close them. However, the process of closure, such as resulting from adding a segment joining the two ends of a polygonal open curve, can introduce entanglements that were not intrinsic to the analysed open curve. Various inventive methods of closure were proposed to minimize the influence of closure on the resulting topology of analysed linear paths (39-44). Unfortunately, in each case, the addition of closing segments changes the geometry of the analysed path in space. This makes that the resulting knot type is not only conditioned by the geometry of the analysed open curve but also by the geometry of somewhat arbitrarily chosen construction of the closing part.

To reveal the intrinsic topology of analysed open curves not affected by a particular set of closing segments a technique involving, multiple, stochastic closures were introduced (45,46). In multiple closure approach, where each closure traces a different path, the distorting effects of individual placements of closing segments are averaged out. One method of multiple, stochastic closure of open curves consists of their closure at “infinity” along a number of equally redistributed directions radiating from a given point (31). In practice, this method corresponds to adding to each of two ends of the open curve a long segment that is parallel to the other added segment. Once the added segments extend beyond the initial open curve their ends are joined by a third segment (see Fig. 2A). The formed closed curve is then characterized with respect to its topology. The closing procedure is then repeated many times, but each time the two long, added segments follow a different direction among many equally redistributed directions radiating from a given point (see Fig 2A). In the analysis reported here, we were closing each of analysed polygonal curves along 100 equally redistributed directions. Importantly, not all directions of closure of a given open curve result in formation of the same knot type and frequently many different knot types are formed when the added parallel segments scan 100 different directions of closure (see Fig. 2B). In the case of the tracing IMR90_chr21-28-30Mb.csv chr_idx:1, the 9_30_ knot was observed most frequently among knots resulting from different directions of closure (see Fig. 2B) and therefore knot 9_30_ is assumed here to represent most closely the topology of the deposited tracing that is shown in Figure 1.

**Figure 2.**
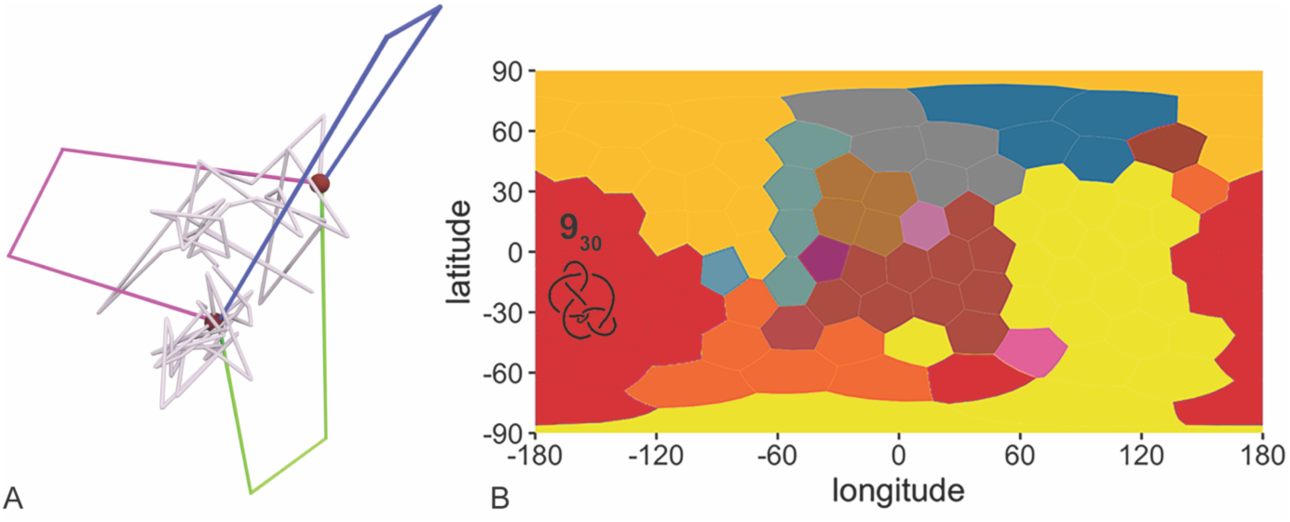
Multiple closure approach reveals the intrinsic topology of an open curve. **A**. Pairs of long segments, that in each case were parallel to each other, were added to both ends of the analysed tracing. Once the added segments extended beyond the analysed polygonal chain they were connected with an additional segment thus forming a closed curve, which knot type can be formally characterized. Different colours of added segments show individual closures. Each analysed tracing was closed 100 times where the directions of added parallel segments were uniformly distributed in the unit sphere. **B**. 100 directions of closures and the resulting knot types. A map representation showing how the closure direction affects the formed knot type. Different colours of Voronoi cells indicate different knot types resulting from the closures in which the added parallel segments were oriented along indicated latitude and longitude angle. In the shown case, the most frequently observed (23%) knot type was 9_30_ knot, although several other individual knot types were also observed in slightly smaller fractions of closures. Notice though that the shown graphs/maps do not preserve the surface of the spheres they represent and the area of individual Voronoi cells, corresponding to given direction of closure, increases as the latitude angle increases. Notice also that shown maps are periodic in horizontal direction and therefore the knot territories adjacent to the left and right border of the map are in fact contiguous with each other. Knot type notation of the most frequently observed knot type is indicated on corresponding “knot countries”. In addition, the diagram of the most frequently observed knot type is also shown. In the matrix shown in panel B and in matrices shown in Figures 3-6, the red colour is used to entries corresponding to most frequently observed knot type conditioned by a given polygonal curve.

We extended then this type of topological analysis to thousands of deposited tracings of the same 2Mb-large chromosome regions of chromosome 21 but acquired in parallel in several thousands of Oligopaint-stained IMR90 cells. Table 1 presents the statistics of the observed knot types. Interestingly, with respect to their topology the deposited tracings behaved similarly to random walks with a moderate size, where the frequency of forming a given knot is inversely correlated with its complexity (47-51). The majority of analysed tracings were unknotted i.e. they formed a trivial knot that has the Alexander-Briggs notation 0_1_. Among non-trivial knots, trefoil knots (3_1_) were most frequently observed. These were followed then by so called figure eight knots that are only knots having 4 crossings (4_1_) and these were followed by two knots with 5 crossings, where twist knots 5_2_ were more frequently observed than the torus knots 5_1_. That predominance of five crossing twist knots over five crossing torus knots was previously observed in studies of random knots (50,51). Among composite knots with two component knots the most frequent are those composed of two trefoils 3_1_#3_1_ and these are followed by these composed of one trefoil knot and one figure-of-eight knot 3_1_#4_1_ which in turn are followed by 3_1_#5_2_ as could be expected from the probability of formation of corresponding prime knots. Among composite knots with three component knots the most frequent are these where each component was a trefoil knot 3_1_#3_1_#3_1_.

**Table 1.**
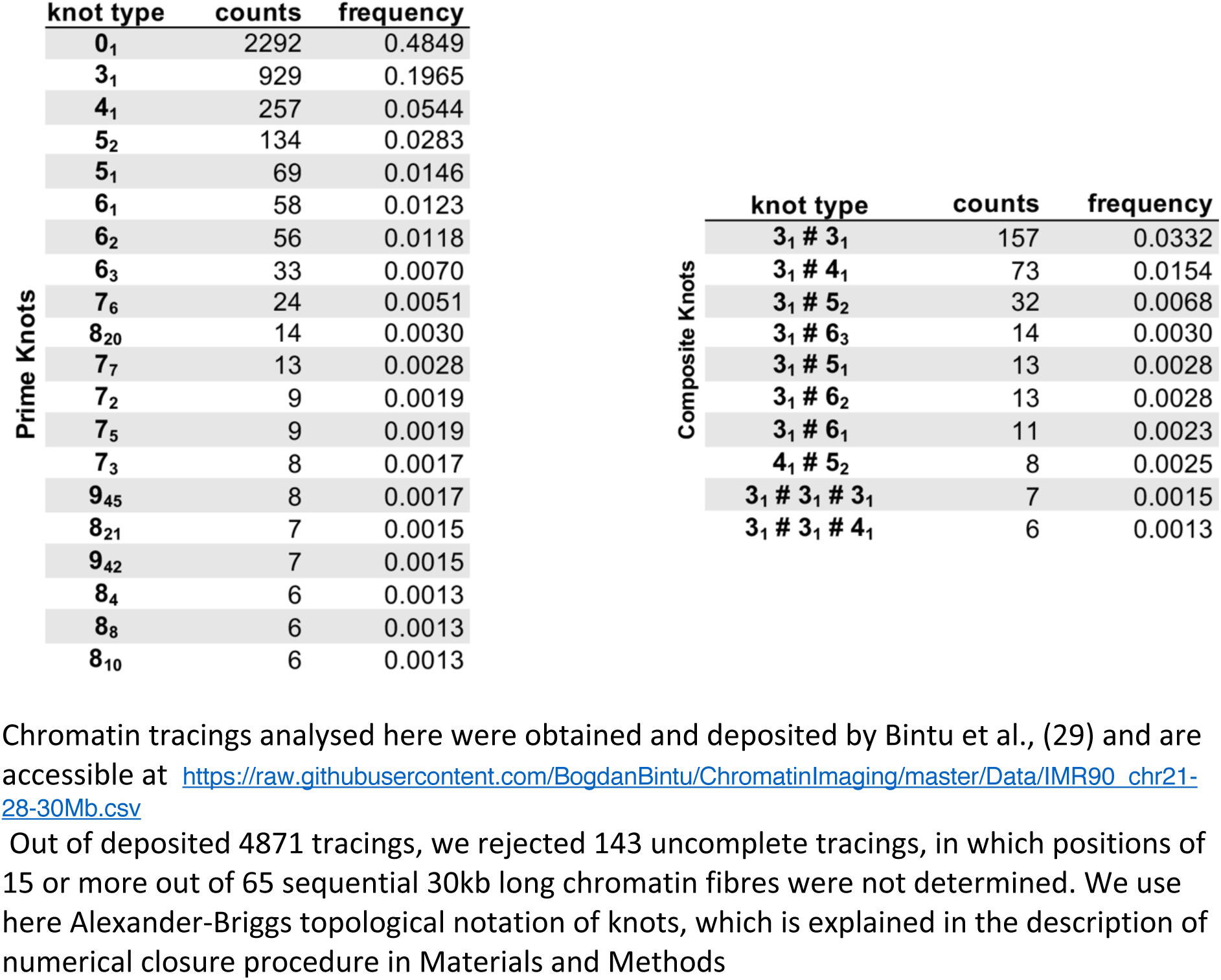
Topological analysis of 4727 deposited chromatin tracings of 2Mb large chromosomal region of chromosome 21 in thousands of cultured human cells.

### Testing the authenticity of detected non-trivial knots

Although nearly half of the analysed chromosome tracings were knotted (see Table 1), the question arises of whether these knots are genuine or are the result of the limited precision in the determination of the path of crowded chromatin fibres. Bintu et al., estimated that the error of determination of the centroid position of each 30 kb-large portion of chromatin fibre is of about 50 nm (29). The diameter of chromatin fibre is of about 10 nm (38) and therefore 50 nm error in the determination of paths of crowded chromatin fibres may result in polygonal tracings that are knotted despite the fact that the traced chromatin was in fact unknotted. Also, the opposite could have happened.

To consider whether the knots reported in Table 1 are genuine or not, let us first perform a thought experiment. Let us assume that we trace highly-crowded polymeric chains using a method that gives us an error of tracing that is larger than many shortest distances between approaching each other non-sequential regions of the traced polymeric chain. Let us consider now that the crowded polymeric chain that we trace is in fact unknotted. A little thought tells us that under such circumstances the produced tracing is very likely to be knotted despite the fact that the traced chain was unknotted. Let us consider now that the crowded polymeric chain that we trace, is in fact knotted and forms a simple knot. Another little thought tells us now that the produced tracing will likely to be still knotted and possibly even more knotted than before. The chance that an error of tracing of crowded polymeric chain that is knotted would produce an unknot is small. The much larger chance that an error prone tracing of a crowded paths will introduce a knot rather than remove a knot has very simple explanation. There is an infinite number of various knots and just one trivial knot. Therefore, if we cram into a pocket our earphone set, it is very likely that it will get knotted. Such a knotted earphone set will not be likely though to get unknotted if we cram it again in a pocket. However, from time to time this may also happen

After this thought experiment, we decided to test numerically the effect of tracing errors on the resulting topology of produced tracings. We first took the same chromatin tracing that is shown in Figure 1 and which multiple closure analysis is shown in Figure 2. We took the original tracing and introduced error-mimicking perturbations, i.e. we displaced by up to 50 nm the position of every vertex, as described in the Methods section. We performed this procedure 10 times, where each time the error-induced perturbations were not correlated with each other but applied randomly. In our analysis of error effects on the detection of chromatin knots, we were inspired by somewhat similar analysis of 10 independent chromosome structure reconstructions from the same single cell Hi-C data (24). For each of 10 error-affected configurations, we performed multiple closure analyses, which results are shown in Figure 3. It is visible that upon perturbations that mimic the intrinsic experimental error, the deposited tracing that was originally forming 9_30_ knot gets very frequently converted into other knots. None of error-perturbed configurations still formed 9_30_ knot. Three of error perturbed configurations formed more complex knots than the unperturbed tracing and were characterized as knots with 10, 11 and more than 12 crossings respectively. Five of error-perturbed configurations formed simpler knots than the unperturbed tracing and resulted in knots with 8, 7, 6, 5 and 4 crossings, respectively. Interestingly, two of error perturbed configurations resulted in a formation of unknotted configuration.

**Figure 3.**
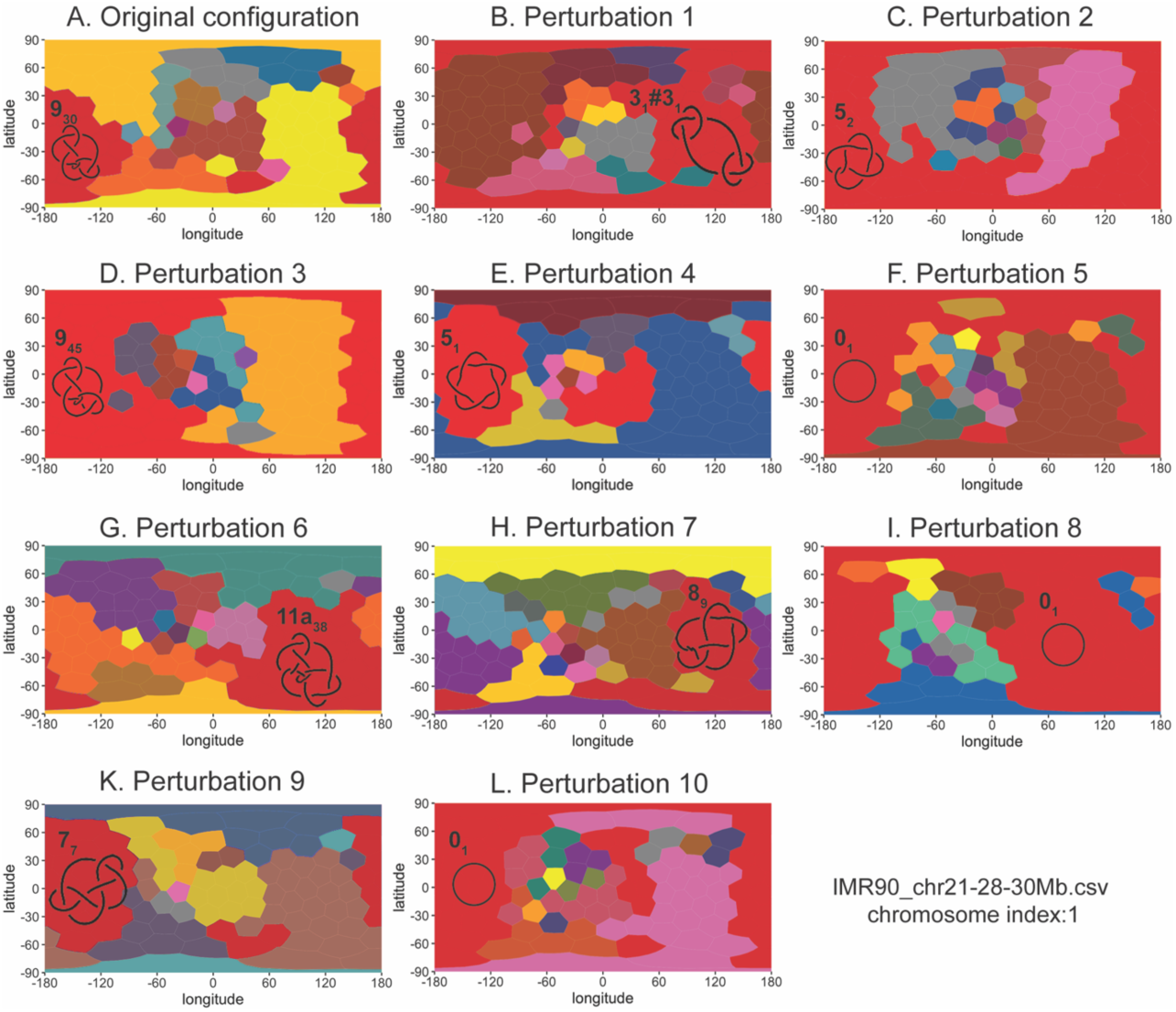
Multiple closure analysis of 10 independently generated error-perturbed configurations of the tracing shown in Figure 1 and analysed in Figure 2. Panel A shows the analysis of the original deposited tracingIMR90_chr21-28-30Mb.csv chr_idx:1. Panels B-L show the closure direction maps for 10 error-perturbed configurations derived from the deposited tracing. Notice that perturbations mimicking the effect of limited experimental precision result in a frequent change of the knot type allocated to a given configurations i.e. the knot type observed for the majority of closure directions. Notice also that three of error perturbed configurations (F, I and L) become unknotted as the majority of their closing directions results in unknots. For each analysed polygonal curve, the knot type notation of the most frequently observed knot type is indicated on corresponding “knot countries”. In addition, the diagram of the most frequently observed knot type is also shown. In each map, the red colour is chosen to indicate Voronoi cells corresponding to directions of closure that result in formation of most frequently observed knot conditioned by a given configuration. The notation and the diagram of the most frequently observed knot, including the trivial knot, is shown in each case.

The analysis presented in Figure 3 indicates that the limited precision of Oligopaint method makes it an unreliable way of determination of chromatin topology for such compact chromatin paths as shown in Figure 1. Our observation that two out of ten error-perturbed configurations resulted in formation of unknotted configurations shows that the deposited knotted tracing is within an error range to a configuration that is unknotted. This latter conclusion is consistent with the notion that traced chromatin fibres may be unknotted and that the knotted character of their tracing is the consequences of the limited precision of the Oligopaint method when applied to crowded chromatin fibres.

### Testing the topological robustness of unknotted tracings

After investigating the effect of experimental errors on the topology of deposited tracings that were classified as knotted, we extended our analysis to tracings that were classified as unknotted. Figure 4 A shows one of the analysed chromosomal tracings that was originally classified as unknotted (IMR90_chr21-28-30Mb.csv chr_idx:209). That tracing is less compact than the knotted tracing shown in Figure 1. For this reason, error-induced perturbations capable of displacing individual vertices by up to 50nm distance are unlikely to produce a knotted path out of such a polygonal curve. Figure 4 B-M show how error induced perturbations affect the topology of this originally unknotted tracing. The original and all 10 of error-disturbed configurations form unknots for the great majority of closure directions. Therefore, the unknotted tracing (IMR90_chr21-28-30Mb.csv chr_idx:209) can be considered as robustly unknotted.

**Figure 4.**
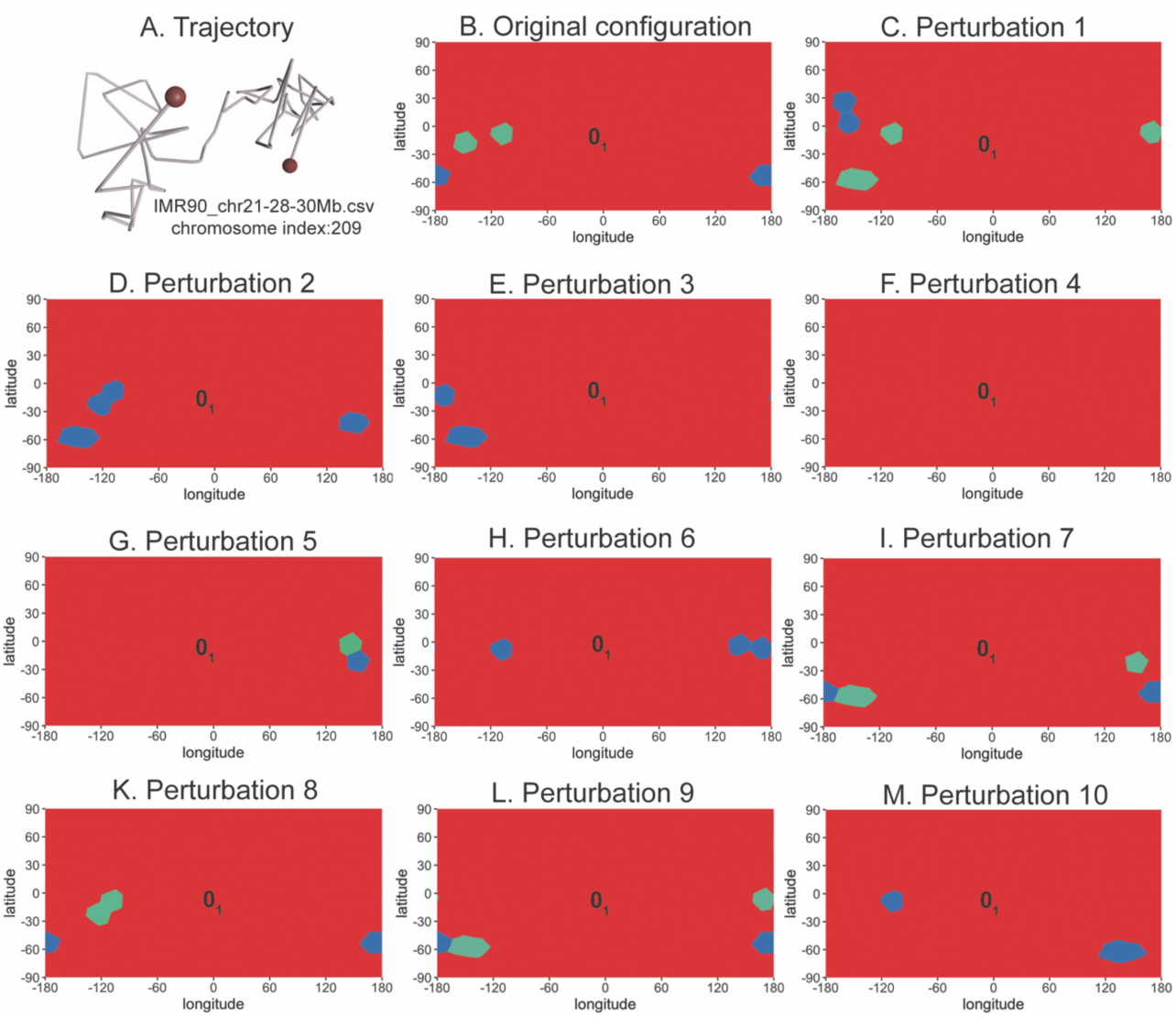
Multiple closure analysis of an unknotted tracing that is robustly unknotted. Panel A shows polygonal chain corresponding to the deposited tracing IMR90_chr21-28-30Mb.csv chr_idx:209. As in Figure 1 and 2, the diameter of the chain is set to correspond to the physical distance of 10 nm. The two beads at the ends have the diameter corresponding to physical distance of 50 nm as this is the reported precision of Oligopaint chromatin tracing. Panels B-L show map representations reporting type of knots resulting from different directions of closure of unperturbed tracing (panel B) and 10 error perturbed configurations derived from unperturbed tracing (panels C-M). Notice that the unperturbed and also all error perturbed configurations are unknotted as the majority of closure directions closes them into unknots.

We extended this type of analysis to all tracings that were classified as unknotted. Among 2292 tracings that were unknotted, 243 were robustly unknotted. Since robustly unknotted tracings are very unlikely to result from an error of tracing method, we can conclude that robustly unknotted tracings report correctly the underlying topology of traced chromatin fibres.

### Search for robust knots

Although we presented earlier arguments that a limited precision of tracing of unknotted but crowded chromatin fibres can easily result in producing knotted tracings, it is also possible that some of traced chromatin fibres were in fact knotted.

We decided therefore to search for robust knots among the analysed tracings. Tracings of robust knots should have the property that the knot type detected in them should resist error induced perturbation i.e. that all of error perturbed tracings derived from a given deposited tracing should show the same knot type. Knotted tracings with such a characteristic would be unlikely to result from experimental imprecisions in tracing procedure. Our search for robustly knotted configurations among 509 tracings classified as trefoil knots was not successful. However, we did find several fairly robust configurations, which we defined as those keeping their original knot type in more than 50% of error-perturbed configurations. Figure 5 shows one of these fairly robust tracing forming a trefoil knot (3_1_) together with multiple closure analysis of its original tracing and 10 error-perturbed tracings. It is visible that 6 out of 10 error-perturbed configurations maintained the knot type of unperturbed tracing. However, 4 out of 10 error-perturbed configurations converted to unknots. This latter observation shows that even such fairly robust tracings forming a trefoil knot can easily be caused by method-specific errors in tracing of unknotted chromatin fibres.

**Figure 5.**
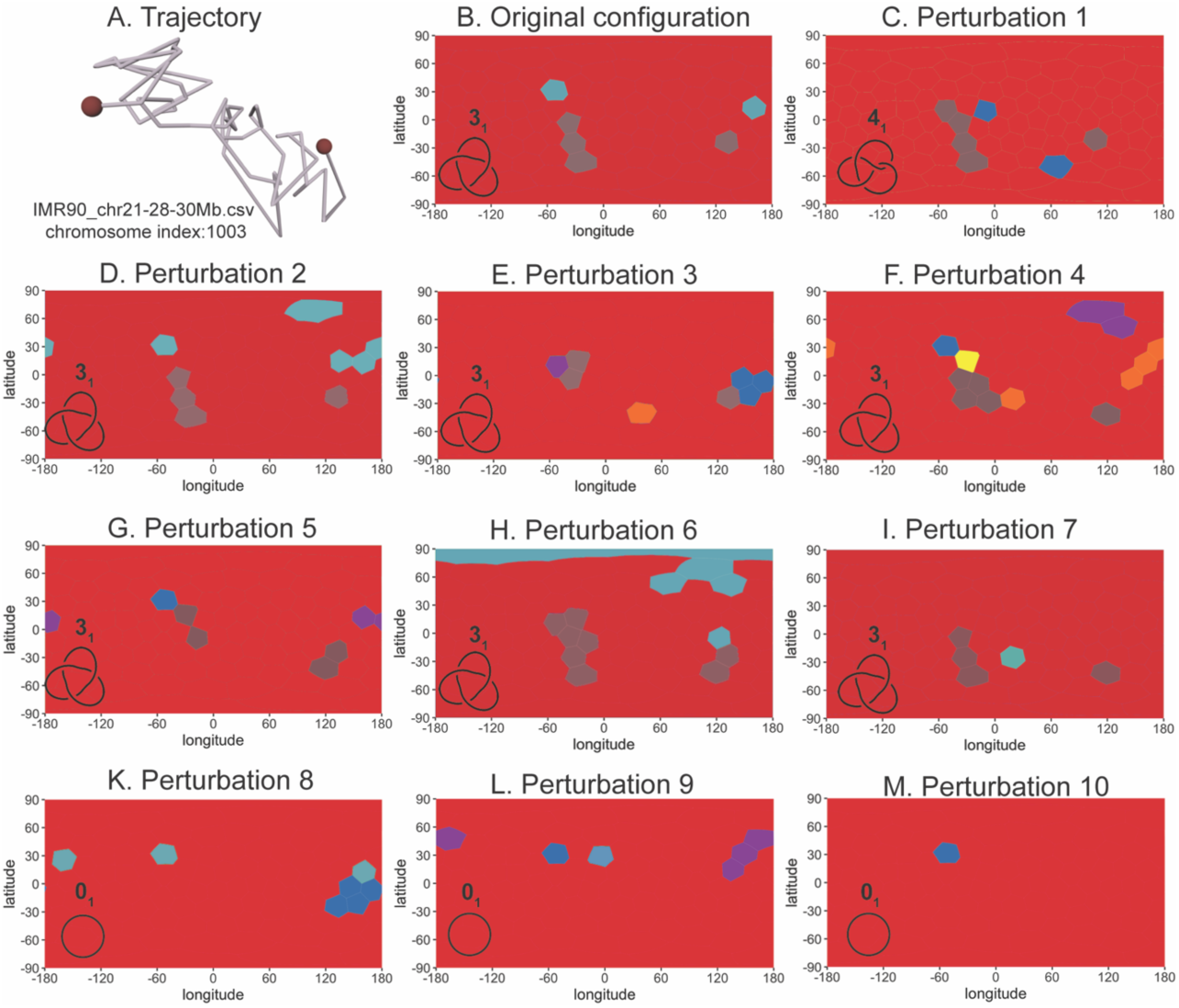
Multiple closure analysis of a fairly robust trefoil-forming tracing. Panel A shows polygonal chain corresponding to the deposited tracing IMR90_chr21-28-30Mb.csv chr_idx:1003. Panels B-M show map representations reporting type of knots resulting from different directions of closure of unperturbed tracing (panel B) and 10 error perturbed configurations derived from unperturbed tracing (panels C-M). Notice that the unperturbed tracing and also 6 of error perturbed configurations are classified as forming 3_1_ knot since the majority of closure directions closes them into 3_1_ knot. However, 4 of the error perturbed tracings are classified as unknotted.

### Error-mimicking perturbations maintain the position of the knotted core as long as the knot is still present

To provide more insights into the question of whether some of the deposited tracings characterized as fairly robustly knotted, report correctly the topology of traced chromatin fibres, we analysed the position of the knotted core in these configurations. The matrix presented in panel A of Figure 6 reports the topology of every subchain of the polygonal chain shown in Fig. 5A. The colour of each cell tells us what is the knot type of the subchain that starts with the vertex indicated on the X axis and ends with the vertex indicated on the Y axis (43). The colour of the cell in the lower left corner of the matrix indicates the knot type of the entire chain and it is 3_1_ knot in this case. We can also see that the smallest subchain that still forms 3_1_ knot and thus forms the knotted core of the chain, starts with vertex 17 and ends with vertex 34. The panels B-L show the topological analysis of subchains in error perturbed tracings that were analysed as entire chains in Figure 5. Interestingly, when error perturbed configurations are still knotted they maintain practically the same position of their knotted core as in the unperturbed configuration. Of course, when error perturbed configurations get unknotted they do not have any more their knotted core. Error-perturbed tracings shown in panels I and K are unknotted when analysed as entire chains, however they form slipknots and therefore some of their subchains form trefoil knots. Panel M shows a superposition of all 10 matrices shown in panels A-L. Such a superposition permits us to visualize more clearly the most persistent patterns.

**Figure 6.**
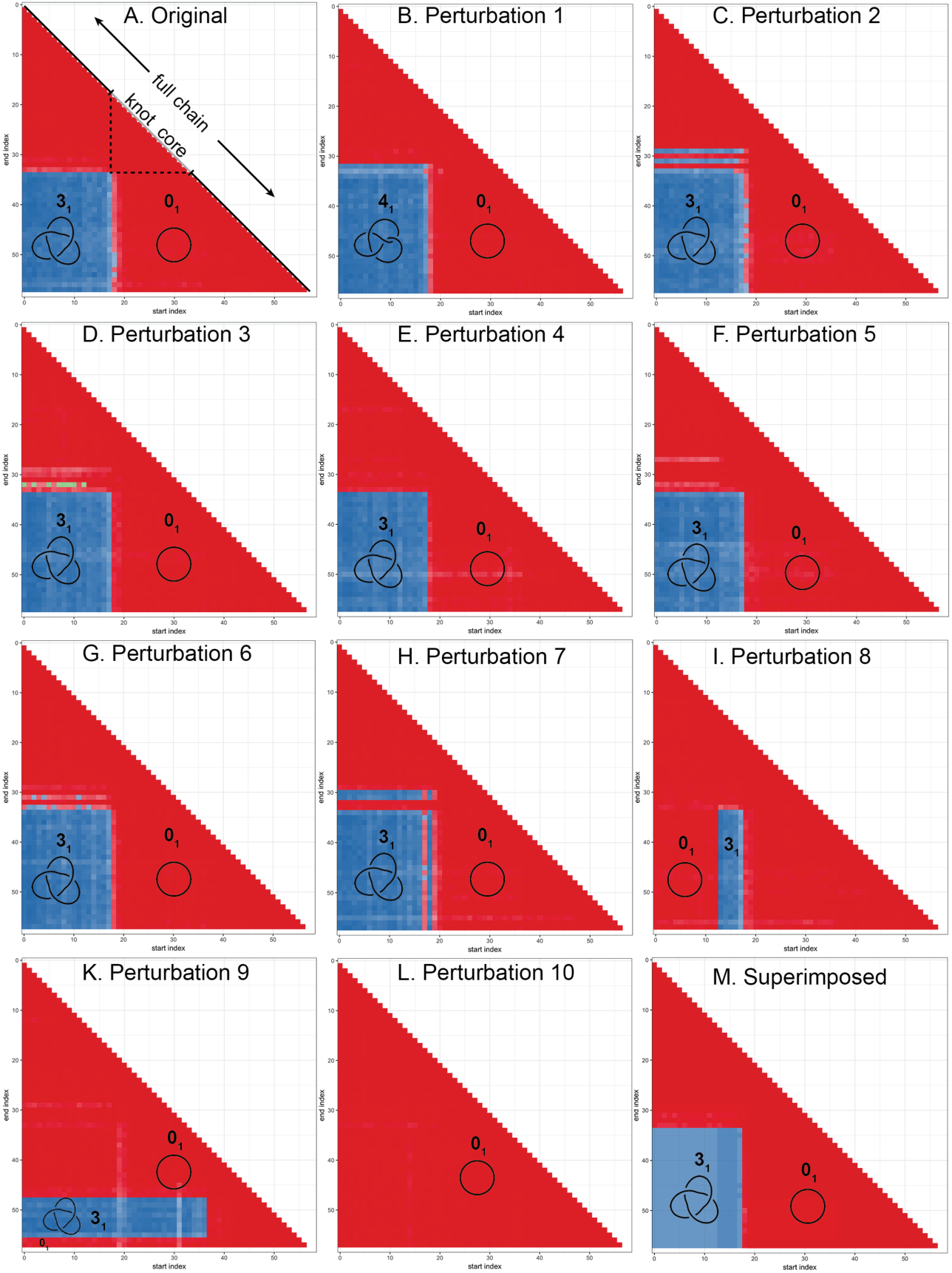
Topological characterization of all subchains in the deposited tracing IMR90_chr21-28-30Mb.csv chr_idx:1003 that forms a fairly-robust trefoil knot (A) and in its error-perturbed configurations (B-M). Colours of individual cells in the matrices A-L indicate the knot type determined by a subchain which starting and ending vertices are indicated on the X and Y axis, respectively. The knot type of each subchain is determined by multiple closure approach presented in Figure 2 but applied to a given subchain. In each matrix the cell in the lower left corner represents the entire chain and its colour indicates the corresponding knot type. Panel A shows that when the deposited tracing is trimmed beyond vertex 18 from the start or beyond the vertex 33 from the end, the truncated tracing becomes unknotted (0_1_). Therefore, the core of the knot in this tracing is located between vertices 18 and 33, as additionally illustrated in panel A. The matrix shown in panel M is a superposition of matrices shown in panels A-L.

The above discussed analysis shows us that if the traced chromatin fibres are indeed knotted, we should be able to determine the position of their knotted core despite the tracing errors.

## Discussion

Strikingly, despite the fact that the majority of analysed by us 4727 deposited chromosome tracings were knotted, it is far from certain that any of the true, underlying, 2Mb long chromatin fibres were indeed knotted. In fact, we found that measurement errors in the chromosome tracing procedure were sufficient to obfuscate the true topology in all deposited tracings that were classified as knotted. In all those tracings, the measured topology was not robust to translation of individual points by distances smaller than the reported error (50 nm) (29). When traced chromatin fibres are highly crowded, as it is frequently the case of interphase chromosomes, the errors of this magnitude can easily produce knotted tracings even if the true trajectory of traced chromatin fibres were in fact unknotted. Presumably for this reason, Bintu et al. (29) did not analyse the topology of chromatin tracings deposited by them.

However, it is also possible that analysed chromatin fibres were in fact knotted. To resolve this ambiguity, we searched for arguments that could tell us whether the observed knots were genuine or artefactual.

The observed spectrum of knots resembles very much the simulated spectra of knotting resulting from topological equilibration that would be expected to occur in crowded chromatin *in vivo*, if topoisomerases were able permit free passages of chromatin fibres through each other in living cells (51). However, essentially the same spectrum of knots would be expected if errors of tracings were “transforming” unknotted but compact chromatin paths into knotted paths of their tracings.

An argument that may be interpreted as pointing towards the notion that *in vivo* chromatin is mainly unknotted was provided though by the absence of robustly knotted tracings i.e. tracings where non-consecutive segments do not approach each other over a distance smaller than 50 nm. Presence of such tracings would provide very strong arguments for the presence of knots in chromatin as such tracings would be very unlikely to result from errors in determination of spatial position of sequentially labelled chromatin portions.

We detected though 243 tracings that were robustly unknotted i.e. tracings that were very unlikely to result from tracing of knotted chromatin fibres. Therefore, in all these cases where tracings had characteristics of correctly reporting the underlying topology of traced chromatin fibres, the analysed 2 Mb-long chromatin fibres were unknotted.

Crucially, when we use experimental data informing us about centroid positions of sequential 30 kb-long chromatin loci within a 2 Mb-long chromosomal region, we can only detect chromatin knots which core length is larger than 150 kb and smaller than 2 Mb. This is because at least five segments of polygonal curve are needed to define a knot upon simple closure (28) and each segment in analysed polygonal tracings corresponds to 30 kb-long chromatin portions. Therefore, if there were chromatin knots with cores smaller than 150 kb, they would not produce knotted parts of polygonal tracings and thus would be missed. However, this compares favourably, though, with single cell Hi-C modelling approach, which due to its 100 kb resolution (24-27), can’t detect chromatin knots if they would be even as large as 500 kb. On the other end of the scale, if the knot core would be larger than 2 Mb (size of analysed fragments), we would also miss it in our analysis. To not miss larger knots, the Oligopaint method would need to be applied to trace chromatin fibres over larger length.

A more definite conclusion about whether chromatin in chromosomes is typically knotted or knot will require improvements in the accuracy of chromosome tracings sufficient to enable topological robustness of the resulting trajectories despite crowded conditions. Until then, the problem will remain knotty.

## ACKNOWLEDGEMENTS

We thank all authors of Bintu et al., (29) for making publicly available all chromatin tracings analysed in this work.

